# Development of constrained peptide inhibitors targeting an oncogenic E3 ubiquitin ligase

**DOI:** 10.1101/2023.04.27.535981

**Authors:** Grasilda Zenkevičiūtė, Wenshu Xu, Jessica Iegre, Hikaru Seki, Yaw Sing Tan, Pamela J.E. Rowling, Fernando Ferrer, Chandra Verma, David R. Spring, Heike Laman, Laura S. Itzhaki

## Abstract

SCF^Skp2/Cks1^ is an E3 ubiquitin ligase, whose substrate specificity is determined by the oncogenic F-box protein Skp2 and the adaptor protein Cks1. A principal target of SCF^Skp2/Cks1^ is the cyclin-dependent kinase inhibitor p27. Elevated levels of Skp2 and reduced levels of p27 are common in a variety of cancers, and there is consequently a need to develop effective inhibitors of the Skp2-p27 interaction. However, conventional small-molecule approaches are challenging due to the extended bi-molecular interface that spans both Skp2 and Cks1, the lack of suitable binding pockets on this surface, and the intrinsically disordered nature of p27. Here, we develop macrocyclic peptides capable of binding to SCF^Skp2/Cks1^ with nanomolar affinities, an enhancement of almost two orders of magnitude over the natural p27 peptide. We show that these macrocyclic peptides inhibit p27 ubiquitination in vitro, restore p27 levels in a breast cancer cell line, and reduce cell proliferation.

## Introduction

The ubiquitin-proteasome system (UPS) is regulated by the sequential action of three enzymes (E1, E2 and E3), of which the E3 ubiquitin ligase confers substrate specificity to the reaction (Fig. 1).

**Figure 1.**
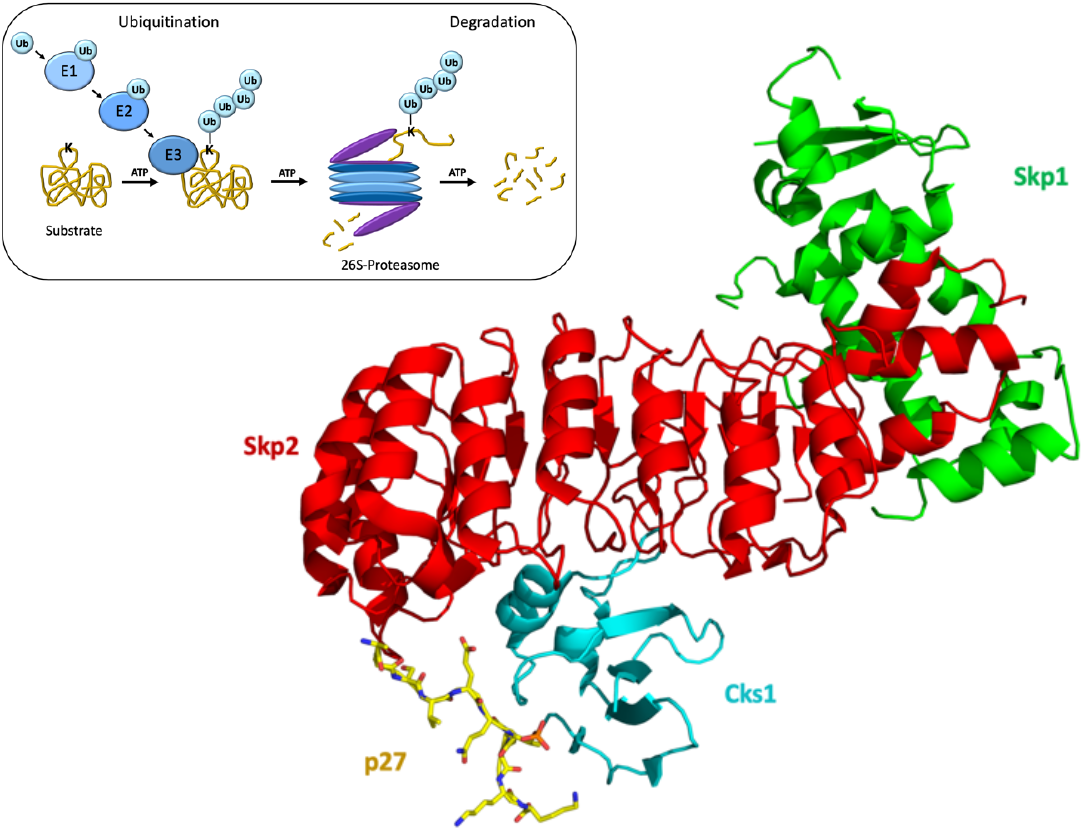
Main: Schematic of the crystal structure of a p27 phosphopeptide bound to Skp2/Cks1 (PDB 2AST) showing that the peptide adopts a turn-like conformation. Inset: Schematic of the ubiquitin-proteasome system.

Following poly-ubiquitination, substrates are recognized and targeted for degradation by the 26S proteasome.^[1]^ The SCF^Skp2^ E3 ligase, a member of the family of SCF ubiquitin ligases, comprises four subunits, Skp1, Cullin1, Rbx1 and F-box protein Skp2, of which Skp2 is the substrate-recognition subunit.^[2]^ A subset of SCF^Skp2^ substrates additionally require the small adapter protein Cks1, which has been shown to form a bimolecular interface together with Skp2 for substrate recognition.^[3]^ Skp2 is overexpressed in a range of cancers, including lymphomas and prostate and breast carcinomas, and is associated with tumor metastasis and poor prognosis.^[4–6]^ Several SCF^Skp2/Cks1^ substrates are tumor suppressor proteins, including the cyclin-dependent kinase 2 inhibitors p21, p27 and p57,^[7]^ degradation of which positively regulates cell cycle progression (Fig. 1).^[8,9]^ Thus, increased levels of Skp2 and reduced levels of p27 are common in many types of cancers,^[10,11]^ and the Skp2-p27 pathway plays a central role in breast cancer initiation and metastasis.^[5]^ There is, consequently, much interest in developing inhibitors of the Skp2/Cks1-p27 interaction. Given the large bimolecular surface involved in binding p27, we chose macrocyclic peptides rather than small molecules for our inhibitor design.

Constrained peptides have been used very successfully to mimic and thereby inhibit protein-protein interactions (PPIs), and some of these peptides are already in clinical trials.^[12–21]^ It is now well understood that constraining peptides in their bioactive helical conformations results in greatly enhanced binding affinities relative to their unconstrained counterparts, and furthermore these so-called ‘stapled’ peptides also possess high stabilities and in some instance cell permeability. There are, however, very few examples of constrained non-helical peptides, in part because the design process is harder than for helical peptides, which can be reliably constrained between i,i+4 or i,i+7 sites with predictable distance requirements.^[22,23]^

Here, we used the natural p27 peptide sequence and the crystal structure of the quaternary Skp1-Skp2-Cks1-p27 complex (Fig. 1B) as the starting point for the design of the peptide inhibitors^[24]^. We employed a two-component double-click chemistry approach for macrocyclization,^[25–27]^ in which the peptide and the linker are independent variables in the design process, thereby enabling the assembly of structurally diverse peptides from a relatively small number of synthesised peptides.

Such diversity is important for finding the best constraint for non-helical peptides, as the design process is not readily predictable in contrast to alpha-helical peptides. The approach yielded macrocyclized peptides with affinities almost 100-fold higher than the linear p27 peptide and capable of inhibiting p27 ubiquitination and degradation and cell proliferation.

## Results

### Peptide and linker design

The phosphorylated p27 peptide that is resolved in the Skp1-Skp2-Cks1-p27 crystal structure is 10 amino acids in length (amino acids 181-190 of p27). Previous solution studies have used much longer 24-residue peptides (175-198 of p27), and a dissociation constant of the order of 7 μM were observed for the interaction with Cks1-Skp2-Skp1.^[24]^ This binding affinity is the same as that of full-length phosphorylated p27.^[24]^ We synthesized the 10-residue peptide, conjugated with a TAMRA fluorophore (TAMRA-Ahx-AGSVEQT(phos)PKK) (Ahx is amino hexanoic acid), and measured its binding affinity for Cks1-Skp2-Skp1 using fluorescence polarisation (FP). The dissociation constant (*K*_d_) was 2.78±0.19 μM, indicating that this short peptide can effectively mimic full-length p27 in its interaction with Cks1-Skp2-Skp1 (Table 1). We further confirmed the binding affinity using ITC (Fig. S1). Next, we use the crystal structure to identify positions on the peptide that are not involved in binding and that could therefore be substituted by unnatural amino acids (UAA) for cross-linking to constrain the peptide in the turn-like conformation that it adopts in its bound form. The UAA sidechain lengths and the linker were chosen based on the distance between the selected sites. Two different pairs of sites were selected for cross-linking that were five or three amino acids apart (Group 1 and Group 2, respectively, in Fig. 2). The sidechains of the UAAs that we incorporated at these sites are of varying length comprising between two and four methylene groups, and all have an azide group at the end to enable macrocyclisation with one of two different alkyne-containing linkers using ‘Click’ chemistry (Fig. 2).^[25,28]^

**Table 1.** Dissociation constants for the binding of macrocyclised p27 peptides to Cks1-Skp2-Skp1 measured by competition FP. The fold increase in binding affinity relative to the linear p27 peptide is also listed. See also Figs. S2 and S3. No binding (N.B.) could be detected for the control peptide in which the three key contacting residues are mutated to Ala. For chemical structures of X1, X2, X3 and X4, see Fig. 2.

| Peptide | Sequence | $K_d$ (nM) | Fold increase |
| --- | --- | --- | --- |
| Control | AGSVAQAPKA | N.B. | N/A |
| Linear | TAMRA-Ahx-AGSVEQT(phos)PKK | $2,777 \pm 190$ | N/A |
| CP1 | AGS(X2)EQT(phos)P(X4)K | $356 \pm 95$ | 8 |
| CP2 | AGS(X2)EQT(phos)P(X3)K | $32 \pm 8$ | 86 |
| CP3 | AGSVE(X3)T(phos)P(X4)K | $81 \pm 4$ | 34 |
| CP4 | AGSVE(X2)T(phos)P(X3)K | $46 \pm 9$ | 60 |
| CP5 | AGSVE(X2)T(phos)P(X2)K | $72 \pm 7$ | 38 |

**Figure 2.**
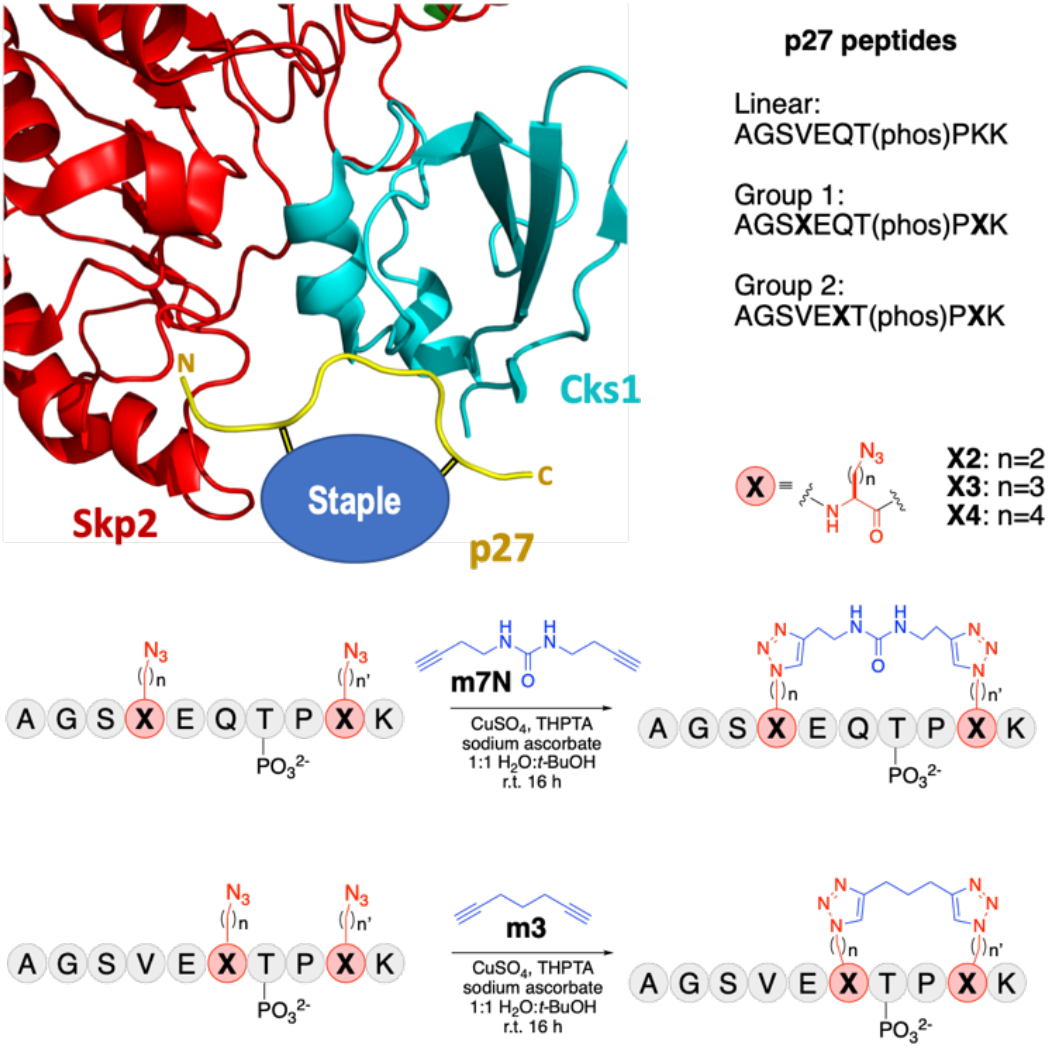
Design of constrained p27 peptides. Two groups of peptides were designed for macrocyclisation at different sites and with different linkers. ‘X’ indicates the positions of the azido-containing unnatural amino acids. Group 1 peptides were constrained with the longer alkyne-containing m7N linker and Group 2 with the shorter alkyne-containing m3 linker. Click chemistry was used to conjugate the azido-containing unnatural amino acids of the peptide to the alkyne-containing linker to constrain it in the bioactive conformation as in the crystal structure. See also Fig. S2.

### Peptide macrocyclization enhances the Skp2-binding affinity by almost two orders of magnitude

To measure the binding affinities of the constrained peptides, we used a competition FP assay, in which the pre-assembled Cks1-Skp2-Skp1 complex was first mixed with a TAMRA-labelled p27 peptide at a concentration such that more than 50% of the binding sites on Cks1-Skp2-Skp1 were occupied. Unlabelled constrained peptides were then titrated into this quaternary complex, and the displacement of the fluorophore-peptide was detected as a decrease in the fluorescence polarization. *K*_d_ values of the constrained peptides (CPs) were obtained by fitting the data as described previously^[29]^ (Fig. 3, Table 1). We also tested a control peptide, in which the three key contacting residues were mutated to Ala (Fig. S4). All but one of the macrocyclised p27 peptides showed 10-fold higher binding affinities than the linear p27 peptide, and one peptide, **CP2**, increased the binding affinity by almost two orders of magnitude. The binding affinity of CP2 was confirmed using ITC (Fig. 3).

**Figure 3.**
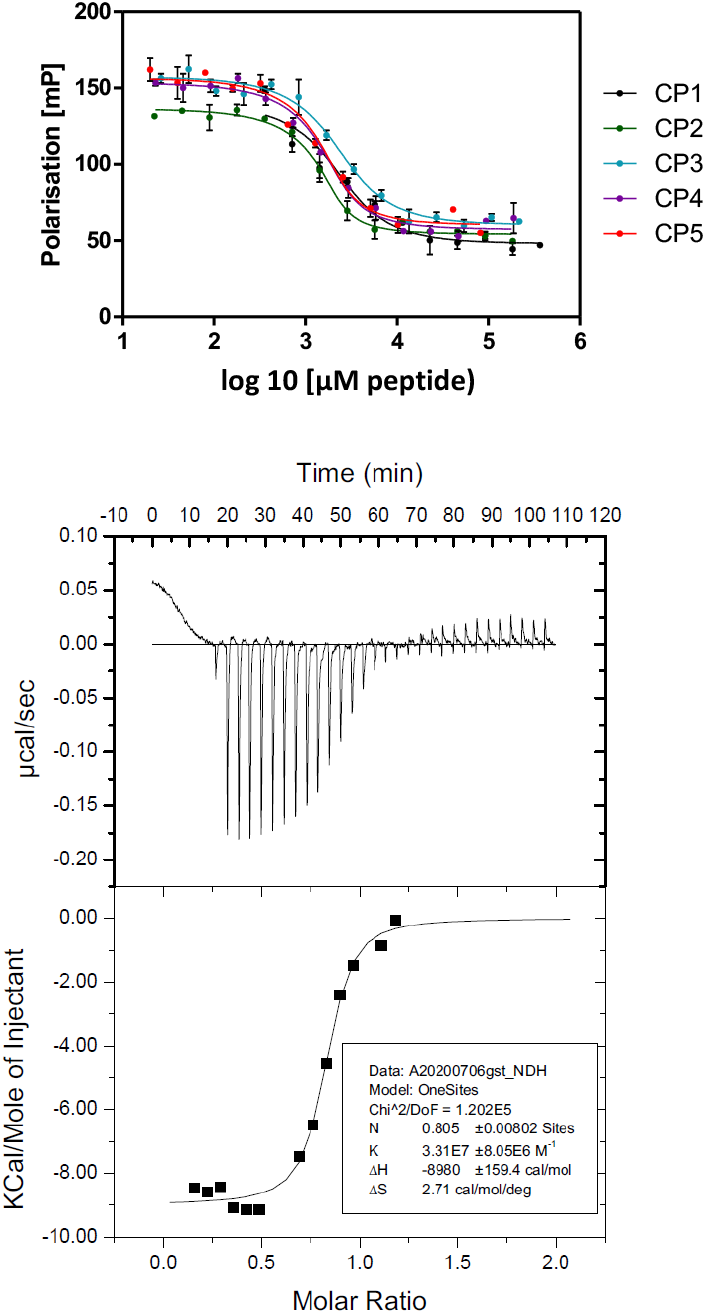
Competition FP analysis of constrained p27 peptides titrated into the complex of Cks1-Skp2-Skp1 and TAMRA-labelled unconstrained p27 peptide. Binding of Cks1-Skp2-Skp1 to macrocyclised peptide CP2 measured by ITC.

### The macrocyclized peptides inhibit SCF^Skp2/Cks1^-mediated ubiquitination of p27

We next investigated the ability of the peptides to inhibit SCF^Skp2/Cks1^-mediated ubiquitination of p27 *in vitro*. In these assays, the peptides were titrated into a reaction mixture containing E1, E2, SCF^Skp2/Cks1^, ubiquitin, ATP and p27 (see also Experimental Section and Figs. S4-S6). Western blot was used to detect poly-ubiquitination of p27 (Fig. 4). In these assays, the addition of the ubiquitin mix, which contains the E1 and E2 enzymes, results in the stochastic mono-ubiquitination of p27 (compare lanes 1 and 2 – see the double band representing unmodified and mono-ubiquitinated p27). This is likely due to proximity-induced ligation of ubiquitin mediated by the E2 enzyme in the ubiquitin mix. This low level of mono-ubiquitination of p27 represents the background level of ubiquitination activity in these assays. However, the addition of the E3 SCF^Skp2/Cks1^ results in massively increased levels of poly-ubiquitinated p27, including a smear and very high molecular weight bands (lane 3), representing longer branched and mixed chains of ubiquitin. Probing for ubiquitin would detect the abundant free 20 mM ubiquitin added into this assay as part of the ubiquitin mix as well as the E1 and E2 enzymes, which are conjugated to ubiquitin as part of the ubiquitin cascade. Another common problem with using antibodies to ubiquitin in Western blots is that weakly associated proteins that are ubiquitinated will be detected. For these reasons we use substrate specific antibodies and conduct experiments under denaturing conditions to ensure that the covalently modified p27 substrate is specifically detected. The results show that all the constrained peptides inhibited p27 poly-ubiquitination to a greater extent than did the linear peptide, consistent with their enhanced Skp2-Cks1-binding affinities.

**Figure 4.**
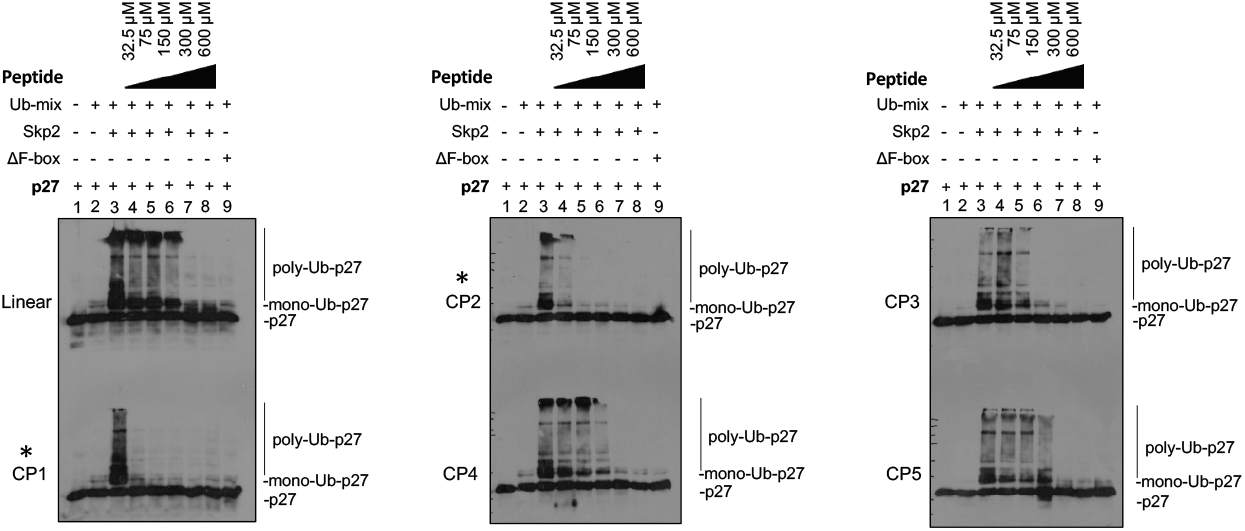
Titration of linear and constrained peptides in the p27 ubiquitination assay. Peptide concentrations are shown on the top of the figure. ‘∆F-box’ corresponds to a truncated variant of Skp2, which lacks the F-box domain and is therefore unable to bind Skp1 and the other SCF subunits resulting in no E3 ligase activity. All experiments were repeated twice, and representative data are shown. The extent of ubiquitination was estimated from the intensity of the poly-ubiquitinated p27 bands. The linear p27 peptide does not achieve full inhibition of p27 ubiquitination even at the highest concentration (600 μM) used. 75 μM of CP1 and CP2 (Group 1 peptides; labelled with *) and 150 μM of CP3 is sufficient for full inhibition. 300 μM of CP4 and CP5 is required for full inhibition.

In addition to the beneficial effect of the chemical constraint on the binding affinities of the peptides, cross-linking should also protect the peptides from proteolysis - an important prerequisite for its use *in vivo*. To this end we measured the protease resistance of the highest affinity macrocyclised peptide, **CP2**, and of the unconstrained version using GluC protease. We also looked at **CP2** fused to a cell-penetrating peptide (CPP); **CP2-CPP** is used in the subsequent cell-based assays (see below). Whereas the majority of linear peptide was degraded within 24 h, almost 90% of stapled peptide **CP2** and **CP2-CPP** remained intact after 24 h of peptidase treatment (Fig. S7).

### The macrocyclized peptides p27 levels in a breast cancer cell line and inhibit cell proliferation

To determine the effects of the peptides in the cell, we used a CPP (sequence RQIKIWFQNRRMKWKK) derived from Antennapedia (Antp) known to facilitate intracellular delivery. Antp was attached at the C-terminus of **CP2**, as the C-terminus is further away than the N-terminus from the Skp2/Cks1-binding site thereby minimising the risk of the Antp peptide affecting the interaction. We also added a 5TAMRA fluorophore for microscopy studies, and we included an amino hexanoic acid (Ahx) linker to provide an adequate spacing between the p27 binding sequence and 5TAMRA dye. The peptide sequence is thus: 5TAMRA-Ahx-AGS-X2-EQT(phos)-X3-KRQIKIWFQNRRMKWKK (**TAMRA-CP2-CPP**). Confocal microscopy of live U2OS cells showed that **TAMRA-CP2-CPP** was able to cross the cell membrane. A proportion of the peptide was localised to the cytosol, as indicated by the diffuse cellular distribution rather than localised endosomal dots (Fig. S8). We then looked at the effects of the peptides on the cellular levels of p27 using the MCF-7 breast cancer cell line. First, we determined whether p27 levels were dependent on Skp2 expression. We found that p27 was almost completely depleted in MCF-7 cells transfected with Skp2 and Cks1 compared with non-transfected cells (compare Lanes 1 and 4 in Figure 5). This result shows that p27 is targeted by SCF^Skp2/Cks1^ for ubiquitination and subsequent degradation. Treatment with proteasome inhibitor MG132 resulted in an increase in p27 levels (compare Lanes 1 and 2), consistent with p27 degradation occurring via the proteasome. We then investigated whether treatment with a constrained p27 peptide could counteract the reduced p27 levels associated with increased Skp2/Cks1 expression. We found that a 16-hour treatment with **CP2-CPP** of cells that had been first transfected with Skp2 and Cks1 restored p27 to the control level (no Skp2/Cks1 transfected) (compare Lanes 3 and 4). This result shows that the constrained p27 peptides have the desired inhibitory effect on the ubiquitination and subsequent degradation of p27 in the cell.

**Fig. 5.**
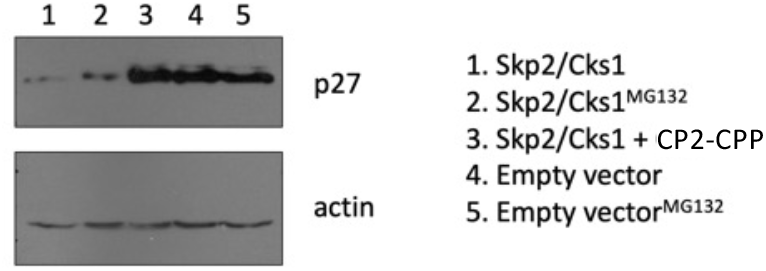
Effect of treatment of Skp2/Cks1-transfected MCF-7 cells with constrained peptide CP2-CPP on the levels of p27. 2.5×10^5^ MCF-7 cells were seeded in 6-well plates and transfected after 24h with either empty vector control or Skp2/Cks1. After another 24 hours, cells were treated with **CP2-CPP** peptide for 16 hours, and the levels of the SCF^Skp2/Cks1^ substrate p27 were then determined by Western blot. Treatment with **CP2-CPP** restores p27 levels to approximately those observed in the control (empty vector transfected) – compare lanes 3 and 4.

We next investigated the effects of the highest affinity peptide **CP2** on cell proliferation. MCF-7 cells were treated with peptides at a concentration of 20 μM for 16 hours and tested using a BrdU (5-bromo-2’-deoxyuridine) incorporation assay measuring DNA replication. BrdU is an analog of the nucleoside thymidine and is incorporated into cellular DNA in place of thymidine during DNA replication. BrdU is detected with primary anti-BrdU antibody, and then HRP-linked secondary antibody is used. Once HRP is activated, BrdU levels can then be determined by measuring the absorbance at 450 nm. Both the CPP that did not have a conjugated CP2 (labelled ‘CPP only’) and the linear peptide conjugated to CPP (labelled ‘linear-CPP’) had only small effects (∼10% decrease) on cell proliferation (Fig. 6). We also tested the control non Skp2-binding peptide (in which the key Skp2-binding residues are mutated to Alanine, see Table 1) conjugated to CPP (labelled ‘Control-CPP’), and this peptide also had only a small inhibitory effect. In contrast, the inhibitory effect was much larger for **CPP-CP2** (35% decrease). We also tested the CPP itself for binding to the Cks1-Skp2-Skp1 complex using ITC, and no binding could be detected (Fig. S9). Thus, the small effect of the CPP on cell proliferation is unlikely to be Skp2-dependent. Since a major function of p27 is as an inhibitor of G1/S phase transition of the cell cycle, the BrdU assay was chosen as a sensitive readout of the effects of any changes in p27 levels during S phase of the cell cycle. In an asynchronous population of cells, only around 30-35% of cells are in S phase. In the 16-hour incubation of MCF-7 cells with **CP2-CPP** (which is less than the duration of one cell cycle), the 35% reduction in DNA replication is both reproducible and 2.5 times greater than the effects of the control peptides. In summary, the results show that constrained p27 peptides can inhibit the ubiquitin ligase activity of the activity of oncogenic SCF^Skp2/Cks1^ and reduce cell proliferation.

**Figure 6.**
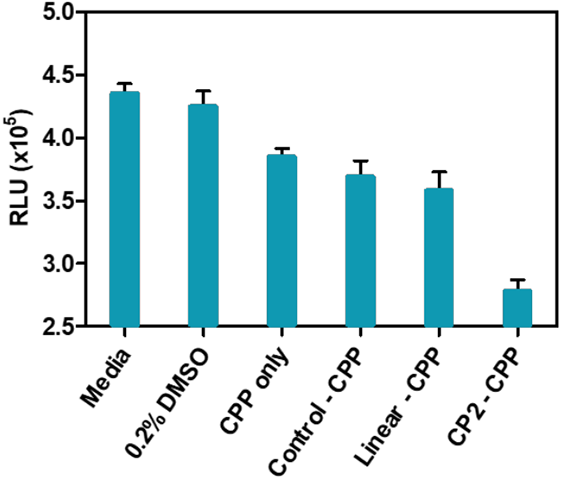
Cell proliferation assay of MCF-7 cells after treatment with CP2-CPP and control peptides. 10^4^ MCF-7 cells were treated with p27 peptides at 20 μM for 16 hours and the BrdU incorporation assay subsequently performed. Error bars are those obtained from triplicate sample measurements.

## Discussion

There are many examples in the literature of constrained alpha-helical peptide inhibitors. However, there are far fewer constrained non-helical peptide inhibitors, and their binding affinities have generally only been marginally improved relative to their linear counterparts.^[22,23]^ In contrast, here we present a striking example in which the constraint of a non-helical peptide induces a dramatic enhancement of binding affinity of almost two orders of magnitude. It is possible that the process of macrocyclisation introduces new binding interactions, for example from the m7N linker or the triazole rings of the peptides effectively creating a triple-constrained peptide structure. But we suggest that the main reason for the dramatic effect of macrocyclisation on binding affinity arises because the bioactive conformation of the p27 peptide is an entropically unfavourable tight turn, which is very effectively induced by the macrocyclic constraint. In contrast, many of the non-helical peptides constrained in previous studies have extended bioactive conformations that are very similar to those sampled when unconstrained and therefore cannot be dramatically improved by constraining them. In summary, the macrocyclic peptides presented here represent a promising new class of inhibitors for E3 ligase SCF^Skp2^, and future work will focus on their further development for use as anti-cancer therapeutics.

### Experimental Section

#### Plasmids

Plasmids containing genes for mammalian and bacterial expression of proteins used in this study are listed in Table S1. Full-length Skp2 and a truncated Skp2 variant lacking the F-box domain that is unable to assemble with the core SCF complex were used.

#### Tissue culture and transfection

HEK 293T, MCF7 and U2OS cells were cultured in Dulbecco’s Modified Eagle’s Medium supplemented with 10% foetal bovine serum and penicillin/streptomycin (Thermo Fisher) at 37 ºC in 5% CO_2_ /95% air atmosphere. HEK 293T cells were seeded in 10 cm tissue culture plates and grown to 50-80% confluency before transfection. PEI (polyethylenimine) or Lipofectamine2000 (Thermo Fisher) transfection reagents were used; DNA was diluted in OptiMEM (serum free media), 3 μl of PEI or Lipofectamine2000 were used per 1 μg of DNA. The mix was vortexed for 10 sec and incubated for 15 min at room temperature prior to drop-wise addition on plates. Plates were incubated at 37 ºC with 5% CO_2/_95% air s for 36-48 hours. For single protein overexpression of Skp2, Skp2ΔFbox-Flag and p27, 3 μg of DNA were transformed per plate.

#### Purification of SCF^Skp2^ complexes from mammalian cells

The SCF^Skp2^ subunits (1.7 μg of each plasmid containing Skp1, myc-Rbx1, Cullin1, and FLAG-Skp2 or FLAG-Skp2ΔFbox) were diluted in Optimen, PEI was added (3 μl per 1 μg DNA), and the DNA was transfected in HEK293T at 50-80% confluency seeded in 10 cm^2^ diameter tissue culture plates. 48 hours post transfection, the cells were harvested in lysis buffer (25 mM Tris-HCl pH 7.5, 225 nM KCl and 1% NP-40) supplemented with protease inhibitors (Sigma-Aldrich) and phosphatase inhibitors (1mM Na_3_VO_4_ and 10 mM NaF). The cell lysate was cleared by centrifugation at 10,600 g for 20 min at 4 ºC. The lysates were incubated with agarose anti-FLAG M2 beads (Sigma-Aldrich) for 4 hours at 4 °C with rocking. Beads were washed with lysis buffer and eluted with FLAG elution buffer (500 μg/mL of FLAG peptide (Sigma), 10 mM HEPES pH 7.9, 225 mM KCl, 1.5 mM MgCl_2_ and 0.1% NP-40) for 2 hours at 4 ºC. Glycerol was added to the final concentration of 15%, and eluates were stored at -20 °C. To evaluate the purification of SCF complexes immunoblotting was performed and probed for anti-FLAG (Sigma), anti-myc (Cell Signalling Technologies), anti-Skp1 (BD Transduction Laboratories) and anti-Cullin1 (Cell Signalling Technologies). The concentration of the SCF protein complexes were determined against known concentrations of BSA standards by Coomassie blue staining. The densitometry of the bands corresponding to Cullin1 and Skp2WT or Skp2ΔFbox was determined using ImageJ (Figs. S6 and S7).

#### Purification of HA-p27 from HEK293T cells

HEK293T at 50-80% confluency seeded in 10 cm^2^ diameter tissue culture plates were transfected with 3 μg of plasmid containing HA-p27. 48 h post-transfection the HEK293T cells were lysed in lysis buffer (25 mM Tris-HCl pH 7.5, 225 mM KCl and 1% NP-40) supplemented with protease inhibitors (Sigma-Aldrich) and phosphatase inhibitors (1mM Na3VO4 and 10 mM NaF). The cell lysate was cleared by centrifugation (14,000 g) at 4 ^◦^C for 20 min. The lysates were incubated with agarose anti-HA beads (Sigma-Aldrich) at 4 ^◦^C for 4 h with rocking. Beads were washed with lysis buffer and eluted with elution buffer (500 μg/mL of HA peptide, 10 mM HEPES pH 7.9, 225 mM KCl, 1.5 mM MgCl2 and 0.1% NP-40) at 4^◦^C for 2 h. Eluates were stored at -20◦C in 5-10 μL aliquots. To evaluate the purification of SCF complexes, immunoblotting was performed and probed for anti-HA (C29F4, Cell Signalling Technologies) secondary anti rabbit Dako.

#### *In vitro* ubiquitination assay

The purified SCF complexes were used at the indicated concentrations in ubiquitination reactions in combination with ubiquitin mix (1X ubiquitination buffer, E1 (100 nM), E2 UBE2DE1 (UbcH5a) (500 nM), ubiquitin (20 mM), Mg-ATP (2 mM)) (Boston Biochem), and p27 was either purified from HEK293T cells or produced by in vitro transcription/translation (IVT), and incubated for 90 min at 30 °C. Proteins were resolved by SDS-PAGE and immunoblotting was performed using anti-HA (Cell Signalling Technologies) or anti-p27 (C-19,Santa Cruz Biotechnologies).

#### Expression in *E. coli* and purification of Cks1 protein and Skp1-Skp2 protein complex

Human Skp2 and Skp1 were cloned as a dicistronic message into the pGEX 4T1 plasmid with the GST– Skp2 fusion coding region preceding that of Skp1 (kindly provided by Brenda Schulman. Full-length Cks1 (1-79) with a N-terminal His-tag was used as described in Seeliger *et al* (2).

Plasmids were transformed into C41 *E. coli* strain (3) and cultured in 5 mL LB medium with antibiotics. The overnight starter culture was then diluted 1:100 in LB medium with antibiotic(s) and grown for 4-5 hours at 37ºC. This culture was used to inoculate 500 mL LB medium with antibiotic(s) (1:100) for large scale expression. Cultures were grown at 37 ºC at 220 rpm until OD600 reached 0.6 and then induced. His-Cks1 expression was induced with 0.2 mM IPTG and incubated at 16 ºC for 18 h. GST-Skp2-Skp1 expression was induced with 0.5 mM IPTG and grown overnight at 25 ºC. Cells were harvested by centrifugation at 4,500 g at 4 ºC for 20 min. The cell pellets were resuspended in a small amount of phosphate buffered saline (PBS) followed by further centrifugation at 4,500 g at 4 ºC for 10 min. Supernatant was discarded and pellets were stored at -20 ºC until needed.

Cell pellets were resuspended in binding buffer, 20 mM Tris-HCl pH 7.5, 150 mM NaCl, 10 mM imidazole, 1 mM DTT (approximately 3 mL of binding buffer per gram of cell pellet) with a protease inhibitor cocktail tablet (Roche) prepared following the manufacturer’s instructions.

Resuspended cells were incubated on ice with frequent shaking for 30 min. Cells were lysed by sonication on ice for a total of 15 mins (30 s on, 30 s off). The cell lysate was cleared by centrifugation (43,150 g, 50 min, 4 ºC). The supernatant, containing soluble proteins, was filtered through a 0.45 μm filter (Sartorius).

For His-tagged Cks1, the cleared lysate was incubated with 1-3 mL of Ni-NTA resin (Qiagen) pre-equilibrated with the binding buffer in a Poly-Prep chromatography column (Bio-Rad) on a rotating wheel for 2 h. The resin was washed with 40 column volumes of binding buffer, and the protein was eluted in 1 mL fractions in elution buffer (20 mM Tris-HCl pH 7.5, 150 mM NaCl, 300 mM imidazole, 1 mM DTT.

For the GST-tagged Skp1-Skp2 heterodimer, the cleared lysate was incubated with 0.5-2 mL of Glutathione Sepharose beads (Cytiva) pre-equilibrated with the binding buffer (20 mM Tris-HCl pH 7.5, 150 mM NaCl, 1 mM DTT) in a Poly-Prep chromatography column (Bio-Rad) on a rotating wheel for 2 h. The resin was washed with 40 column volumes of lysis buffer, and then eluted in 0.5 mL fractions in elution buffer (20 mM Tris-HCl pH 7.5, 150 mM NaCl, 10 mM glutathione, 1 mM DTT).

Size-exclusion chromatography (SEC) was used as a final step to purify both His*-*Cks1 and GST-Skp1-Skp2 complex. The proteins were then injected onto a HiLoad 26/60 Superdex 75 or a HiLoad 16/60 Superdex S75 columns (Cytiva) pre-equilibrated with SEC buffer (20 mM Tris-HCl pH 7.5, 150 mM NaCl, 1 mM DTT). Fractions containing pure protein of interest were pooled and concentrated. Protein concentration was determined using the calculated extinction coefficient at 280 nm (4), His-cks1, 15,930 M^-1^ cm^-1^ and GST-Skp2-Skp1, 96,830 M^-1^ cm^-1^. Proteins were flash-frozen in liquid nitrogen and stored at -80 ºC. Protein purity and identity were confirmed using SDS-PAGE and mass spectrometry.

#### Peptide synthesis and macrocyclisation

The linear p27 peptides containing unnatural amino acids bearing azide functionalities were synthesised by Cambridge Peptides. For the macrocyclization reactions, all solvents were degassed for 1 h prior to use. A solution of CuSO4 (1 equiv.), sodium ascorbate (NaAs, 3 equiv.) and tris(3-hydroxypropyltriazolylmethyl)amine (THPTA, 1 equiv.) in H2O was added to a sealed reaction flask containing diazido-peptide (1 equiv.) and dialkynyl linker (1:1 equiv.) in 1:1 H2O/tBuOH (2 mL). The mixture was stirred at room temperature for 16 h. Progress of the reaction was monitored on LC-MS. Another portion of the linker and catalysts was added if reaction was not completed after 16 h.

#### Peptide purification using HPLC

HPLC was performed on semi-preparative HPLC Agilent 1260 Infinity using a Supelcosil ABZ+PLUS column (250 mm ⇥ 21.2 mm, 5 mm). Samples were run at a flow rate of 20 mL per min at 25 ºC and separated using a linear acetrontirile gradient from 5-50% in MeCN/H_2_O (0.01% TFA). Absorbance was monitored at 254 nm and 220 nm. The masses of the macrocyclised peptides were confirmed by LC-MS (Supplementary Table S2).

#### Fluorescence Polarisation (FP) assays

The proteins were prepared for Direct FP assays by serially diluting 1:2 in SEC buffer containing 0.01 % Tween-20 in triplicate to produce a 12-point curve in a 50 μl volume. TAMRA-labelled peptide (TAMRA-Ahx-AGSVEQT(phos)PKK) was dissolved in water as a stock solution and the concentration determined using NanoDrop2000 (Thermo Scientifc). Absorption was measured at 556 nm and the extinction coefficient of 5-TAMRA of 8900 L mol^-1^ cm^-1^ was used. In each well, TAMRA-labelled peptide (20 nM, 50 μL) was mixed with the serially diluted protein and incubated for 30 min at room temperature before the measurement. Fluorescence polarisation (FP) was measured using a CLARIOstar microplate reader (BMG Labtech) using the following filter settings: excitation filter 540-20 nm; emission filter 590-20 nm, and dichroic mirror LP 566 nm. The data were analysed using GraphPad Prism 5.0 and the dissociation constant, *K*d, was determined using the following equation assuming the ratio between the concentration of the bound and that of the total TAMRA-labelled peptide is proportional to the polarisation change:

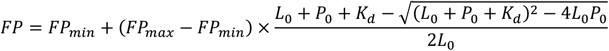

where FP is the fluorescence polarisation, FPmin is the minimum FP, FPmax is the maximum FP, P_0_ is the total concentration of protein, L_0_ is the total concentration of TAMRA-labelled peptide, and *K*_d_ is the dissociation constant.

For the competition FP assays, the highest concentration of the macrocyclic peptide or non-phosphorylated peptide required was prepared up to 5 mM in water, and serially diluted 2-fold for a 12 or 16-point titration curve in triplicate. TAMRA-AGSVEQT(phos)PKK peptide (20 nM) and Cks1-Skp2-Skp1 (4 μM) were incubated in a SEC buffer containing 0.01 % v/v Tween 20. In each well, the unlabelled macrocyclic peptide (up to 5 mM, 50 mL) was mixed with the Cks1-Skp2-Skp1-TAMRA-AGSVEQT(phos)PKK solution (50 mL), and incubated for 30 min at room temperature before the measurement was taken. The data were analysed with GraphPad Prism 5.0 using the equation below, as previously described by Wang. ^[1]^

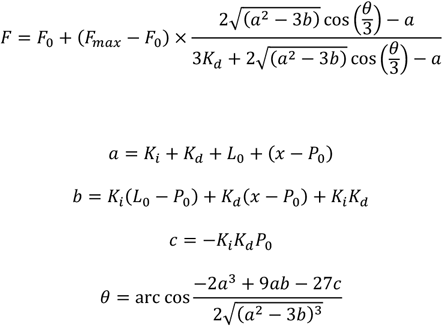

where FP is fluorescence polarisation, *K*_d_ is the dissociation constant determined for the TAMRA-labelled peptide, *K*_i_ is the dissociation constant of the unlabelled macrocyclic peptide, L_0_ is the total concentration of the TAMRA-labelled peptide, P0 is the total concentration of Cks1-Skp2-Skp1 complex, x is the concentration of the unlabelled macrocyclic peptide, FP_0_ is the fluorescence polarisation when no TAMRA-labelled peptide is bound to Cks1-Skp2-Skp1, and FP_max_ is the fluorescence polarisation when all TAMRA-labelled peptide is bound.

#### Isothermal titration calorimetry

The binding of peptides to Skp1-Skp2-Cks1 complex was measured using isothermal titration calorimetry (ITC) using a Microcal VP-ITC (Malvern Panalytical). The protein complex of GST-Skp2-Skp1 and His-Cks1 was preformed first by mixing the proteins in a 1:2 molar ratio in 50 mM Tris-HCl, pH 7.5, 150 mM NaCl and 2 mM DTT. The complex was then run on a size-exclusion chromatography column (Superdex™ 200 Increase 10/300 GL, Cytiva) to remove excess Cks1. The concentration of the complex was determined using an extinction coefficient of 112,760 M^-1^ cm^-1^. The titration was performed at 10 °C by an initial injection of 3 μL of peptide (3 μM) followed by 46 injection steps at 6 μM into GST-Skp2-Skp1-His-Cks1 (4 uM). The data was analysed with MicroCal Origin software.

#### Protease digestion experiments

Endoproteinase GluC (50 ng, New England BioLabs) and caffeine (20 μl of a 7 mg/ml stock in water (to act as an internal control)) was added to p27 peptide (either linear, stapled CP2 or CP2 tagged with CPP) (400 μM) in endoproteinase GluC enzyme buffer (New England BioLabs), and caffeine as internal standard (20 μL of a 7mg/mL stock solution in water), and the volume was made up to 50 μL with sterile water. The reaction was incubated with shaking at 550 rpm at 37 °C. At each time point, a 5 μL aliquot was taken and centrifuged at 10,000 rpm at 4 °C, and the supernatant was diluted 4 times with water before being loaded onto an analytical HPLC (Agilent 1260 Infinity, Supelcosil ABZ+PLUS column (150 mm × 4.6 mm, 3 μm)) and eluted with a linear gradient system (solvent A: 0.05% (v/v) TFA in water; solvent B: 0.05% (v/v) TFA in acetonitrile). The data shown are representative of one biological experiment performed in triplicate.

## Supporting Information

Table S1. Plasmids used in this study.

Table S2: LC-MS data of macrocyclised peptides after the click reaction.

Figure S1. Binding of Cks1-Skp2-Skp1 to a 10-residue phosphorylated p27 peptide by ITC. Figure S2. Chemical structures of constrained p27 peptides.

Figure S3. Competition FP and ITC experiments of binding of Cks1-Skp2-Skp1 to a control p27 peptide. (AGSVAQAPKA), in which all three key contacting residues were mutated to alanine.

Figure S4. Immunoprecipitation of SCF^Skp2 WT^ and SCF^Skp2−Fbox^.

Figure S5. SDS-PAGE of immunoprecipitation of SCF^Skp2 WT^ and SCF^Skp2−Fbox^.

Figure S6. *In vitro* p27 ubiquitination assay performed using p27 substrate prepared by immunoprecipitation from HEK293T cells.

Figure S7. Timecourse of proteolytic degradation by GluC protease of linear p27 peptide, CP2, and CP2-CPP.

Figure S8. Live U2OS cells were treated for 4 hours with TAMRA-CP2-CPP at 20 μM prior imaging. Figure S9. ITC of the 16aa cell-penetrating peptide Antp (sequence RQIKIWFQNRRMKWKK) titrated into the Skp2-Skp1-Cks1 complex.

## Acknowledgements

GZ was supported by a Cancer Research UK Cambridge Cancer Centre PhD studentship. LSI acknowledges the support of a Senior Fellowship from the UK Medical Research Foundation and grants into her lab from the Pancreatic Cancer Research Fund and CRUK (C17838/A22676 and C17838/A27225). GZ acknowledges the support of a Fieldwork Fund grant from the University of Cambridge School of the Biological Sciences. YST and CSV thank A*STAR (IAF-PP H17/01/a0/010) for support.

## Notes

### Competing Interest Statement

The authors have declared no competing interest.

