## Supporting Information for "Development of constrained peptide inhibitors targeting an oncogenic E3 ubiquitin ligase"

**Abstract:** *$SCF^{Skp2/Cks1}$ is an E3 ubiquitin ligase, whose substrate specificity is determined by the oncogenic F-box protein Skp2 and the adaptor protein Cks1. A principal target of $SCF^{Skp2/Cks1}$ is the cyclin-dependent kinase inhibitor p27. Elevated levels of Skp2 and reduced levels of p27 are common in a variety of cancers, and there is consequently a need to develop effective inhibitors of the Skp2-p27 interaction. However, conventional small-molecule approaches are challenging due to the extended bi-molecular interface that spans both Skp2 and Cks1, the lack of suitable binding pockets on this surface, and the intrinsically disordered nature of p27. Here, we develop macrocyclic peptides capable of binding to $SCF^{Skp2/Cks1}$ with nanomolar affinities, an enhancement of almost two orders of magnitude over the natural p27 peptide. We show that these macrocyclic peptides inhibit p27 ubiquitination in vitro, restore p27 levels in a breast cancer cell line, and reduce cell proliferation.*

**Table S1. Plasmids used in this study**

| Construct | Vector | Residue numbers (full length unless indicated otherwise) |
|---|---|---|
| Skp2WT-Flag | pcDNA3 | |
| Skp2ΔFbox-Flag | pcDNA3 | Δ93-144 |
| p27-HA | pcDNA3 | |
| Skp1 | pcDNA3 | |
| Cullin 1 | pcDNA3 | |
| Myc-Rbx1 | pcDNA3 | |
| GST-Skp2-Skp1 | pGEX4T1 | Skp2 101-436<br>Skp1 1-37, 44-70, 83-162 |
| His-Cks1 | pRSETA | |

**Table S2: LC-MS data of macrocyclised peptides after the click reaction.** Calculated m/z ratios are for [M+2H]2+.

| Peptide | Linker | m/z calculated | m/z observed |
|---|---|---|---|
| CP1 | m7N | 692.7 | 692.3 |
| CP2 | m7N | 685.7 | 685.4 |
| CP3 | m3 | 649.2 | 648.9 |
| CP4 | m3 | 635.3 | 634.9 |
| CP5 | m3 | 628.3 | 627.9 |

**Figure S1. Binding of Cks1-Skp2-Skp1 to a 10-residue phosphorylated p27 peptide by ITC.** Measurements were made at 10 °C.

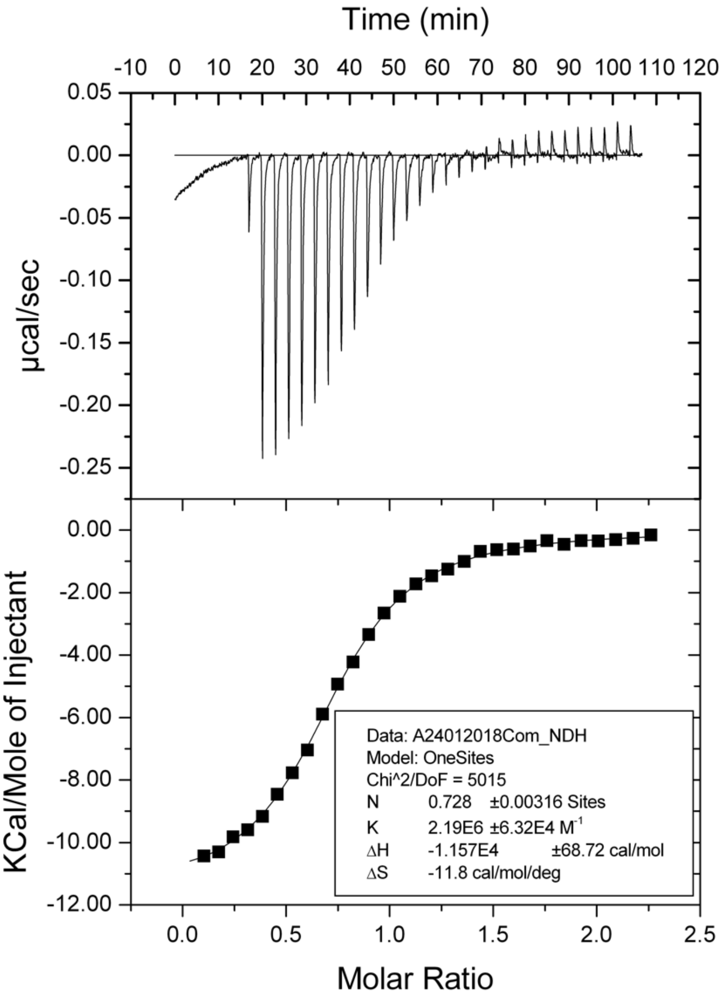

**Figure S2. Chemical structures of constrained p27 peptides.**

**CP1**

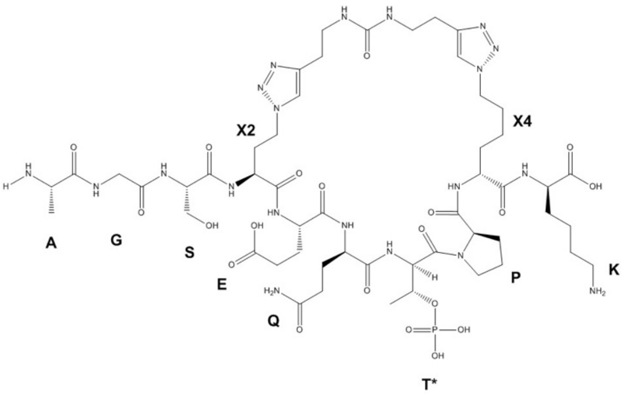

**CP2**

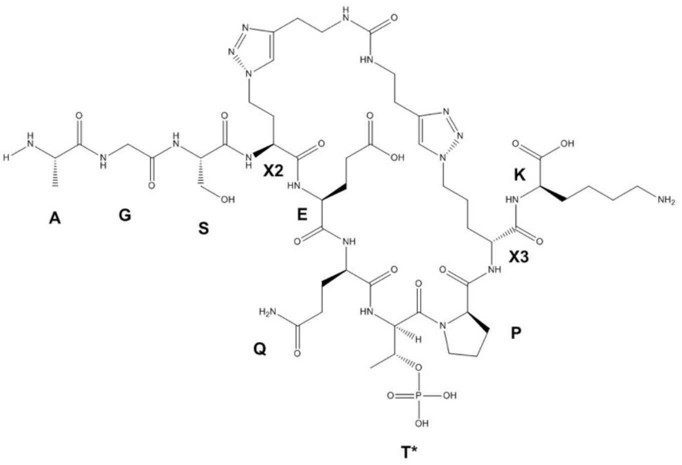

**CP3**

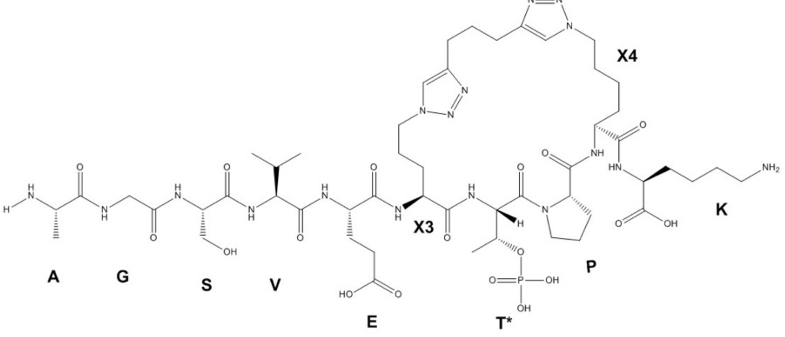

**CP4**

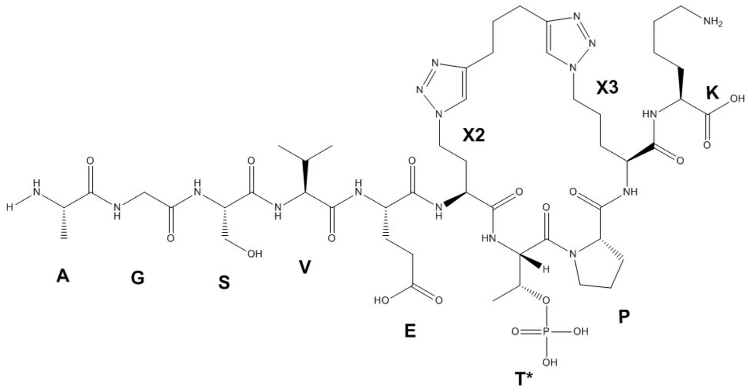
NH2
X3
K
OH
HN
O
X2
N
H
O
A
G
S
V
OH
HO
O
E
T*
P
O=P-OH
OH

**CP5**

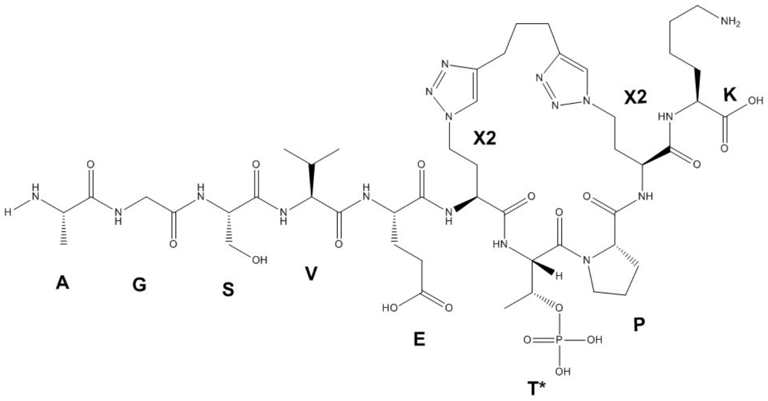
NH2
X2
K
OH
HN
O
X2
A
G
S
V
OH
HO
O
E
T*
P
O=P-OH
OH

**Figure S3. Competition FP and ITC experiments of binding of Cks1-Skp2-Skp1 to a control p27 peptide (AGSVAQAPKA), in which all three key contacting residues were mutated to alanine.**

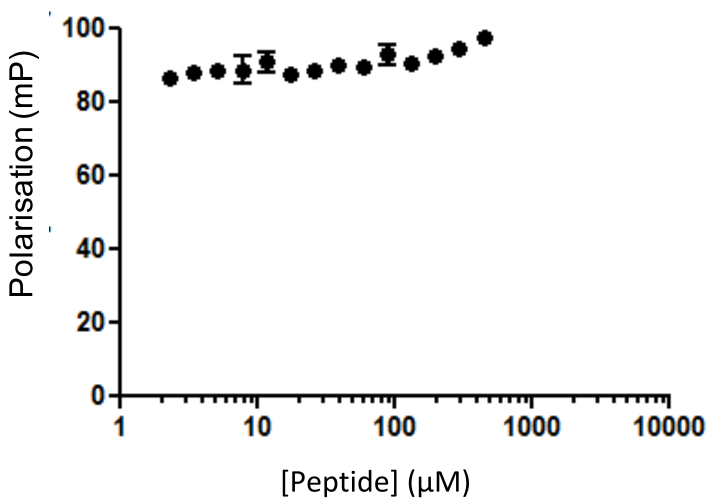


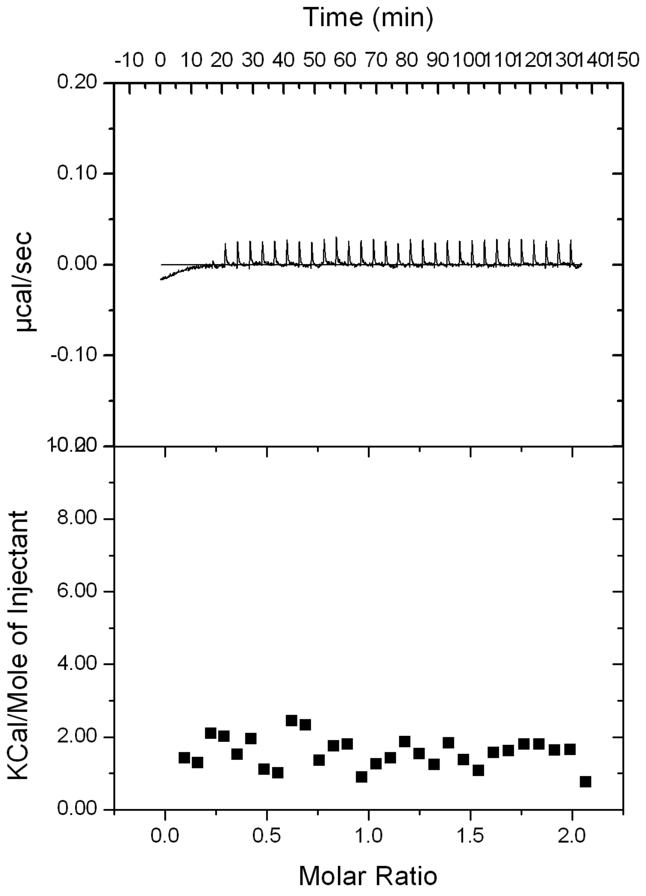

**Figure S4. Immunoprecipitation of SCF$^{Skp2\ WT}$ and SCF$^{Skp2\Delta Fbox}$.** The SCF ligase complex was immunoprecipitated with anti-FLAG beads, and samples were tested for the presence of Skp2, Cullin1, Skp1 and Rbx1 proteins using specific antibodies. Skp1, Cullin1 and Rbx1 are absent in SCF$^{Skp2\Delta Fbox}$ sample, confirming deletion of the Fbox and consequent failure to assemble the E3 complex. Molecular weights are shown in KDa.

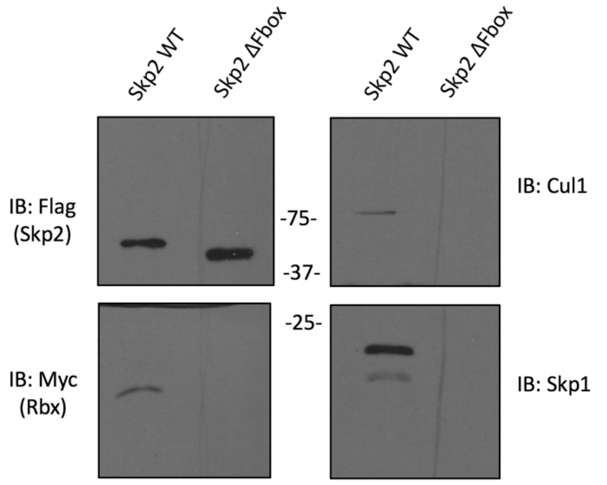


**Figure S5. SDS-PAGE of immunoprecipitation of SCF$^{Skp2\ WT}$ and SCF$^{Skp2\Delta Fbox}$.** The proteins were stained with Coomassie blue, and the bands were quantified using ImageJ. BSA (bovine serum albumin) standards are also shown.

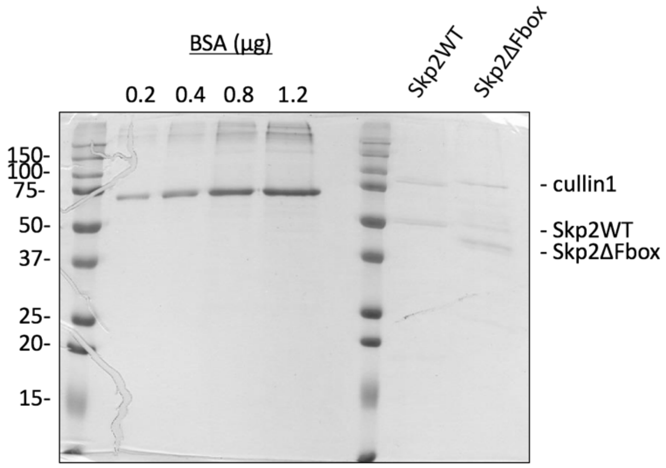

**Figure S6. *In vitro* p27 ubiquitination assay performed using p27 substrate prepared by immunoprecipitation from HEK293T cells.** $SCF^{Skp2}$ was titrated to determine the optimal concentration for p27 ubiquitination.

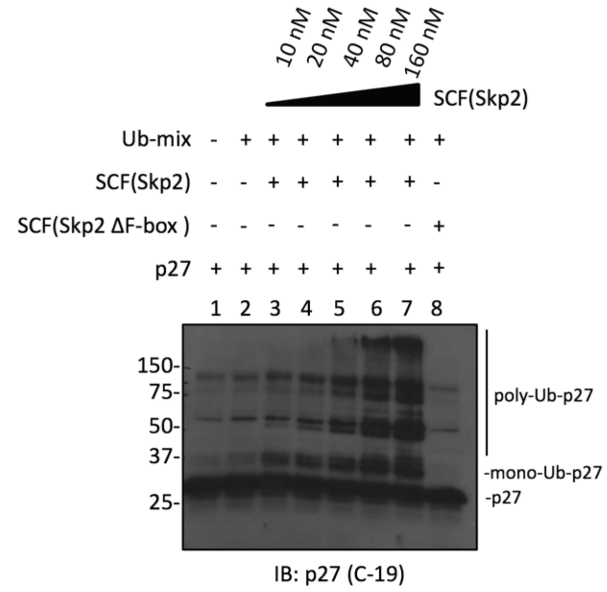

**Figure S7. Timecourse of proteolytic degradation by GluC protease of linear p27 peptide, CP2, and CP2-CPP.** The integral of the elution peak corresponding to the intact peptide was measured against that of caffeine as the internal standard. The majority of linear p27 peptide was degraded within 24 h, whereas almost 90% of stapled CP2 peptide and CP2-CPP remained intact after 24 h of peptidase treatment. Results are shown as mean +/- SD.

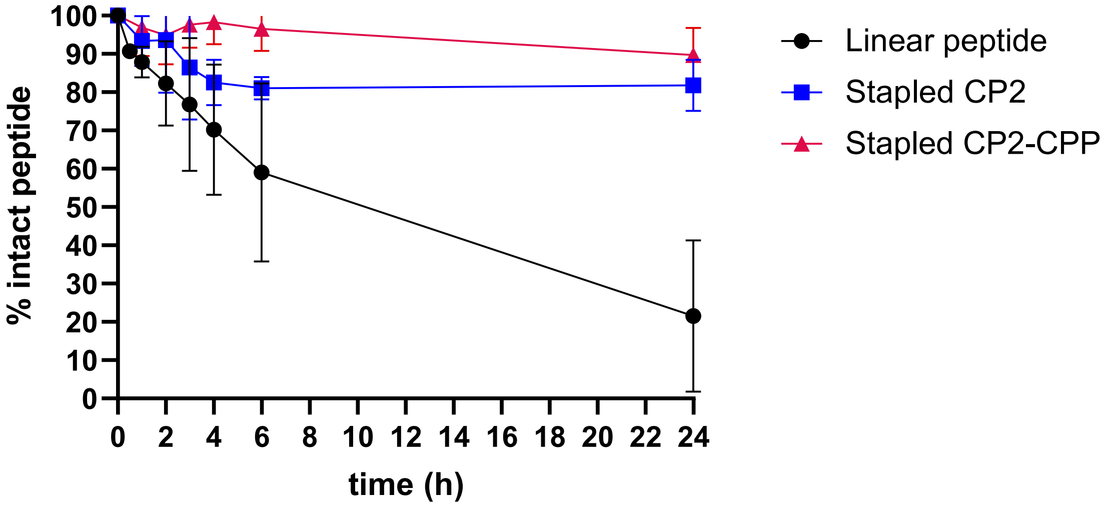

**Figure S8. Live U2OS cells were treated for 4 hours with TAMRA-CP2-CPP at 20 µM prior imaging.** Images were taken on Leica tandem confocal microscope using 40X objective. A. White light. B. Blue dye (DAPI) showing nuclei staining. C. Red corresponds to TAMRA-labelled peptide. D. Overlay.

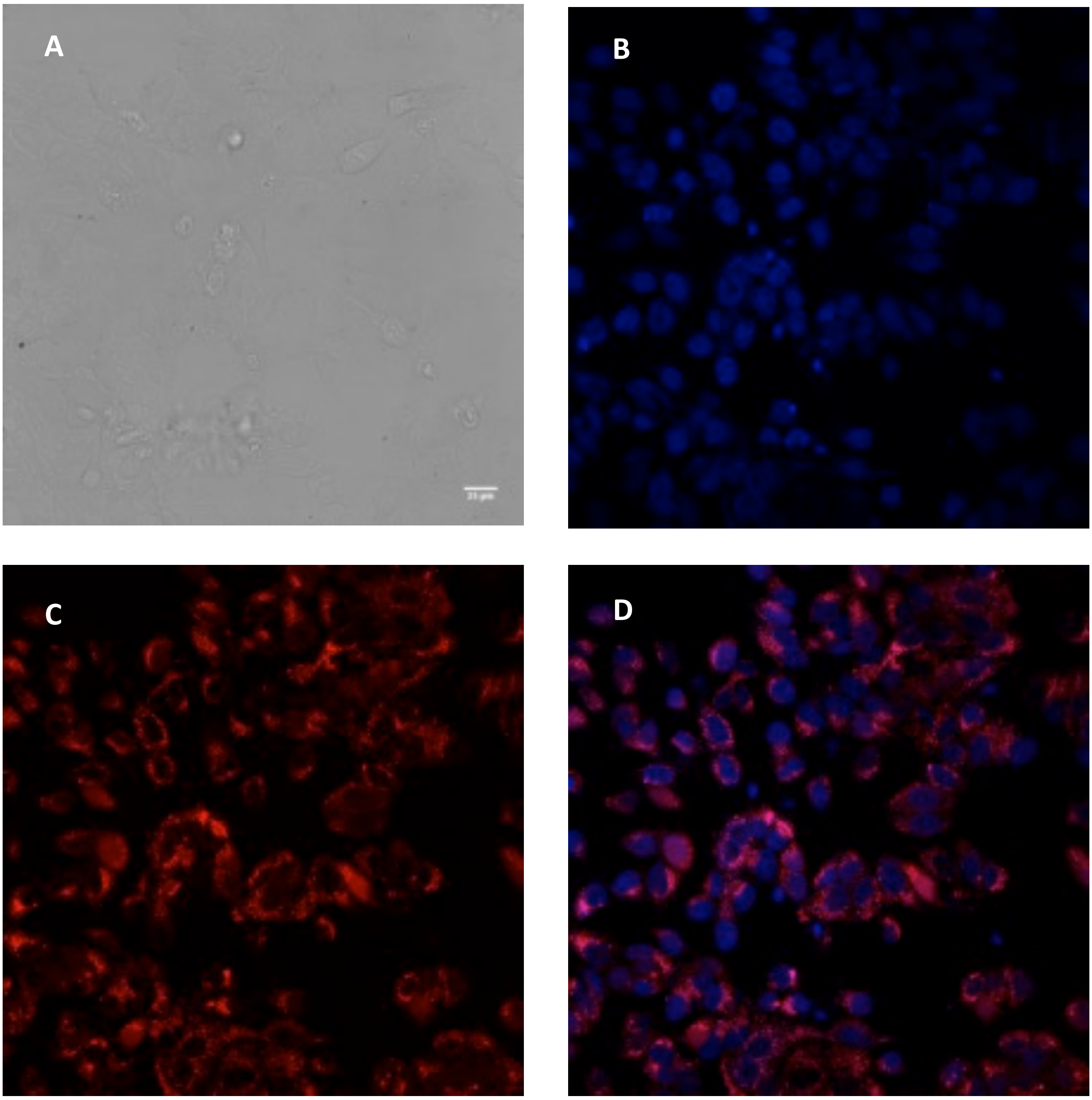

**Figure S9. ITC of the 16aa cell-penetrating peptide Antp (sequence RQIKIWFQNRRMKWKK) titrated into the Skp2-Skp1-Cks1 complex.** Measurements were made at 10 °C.

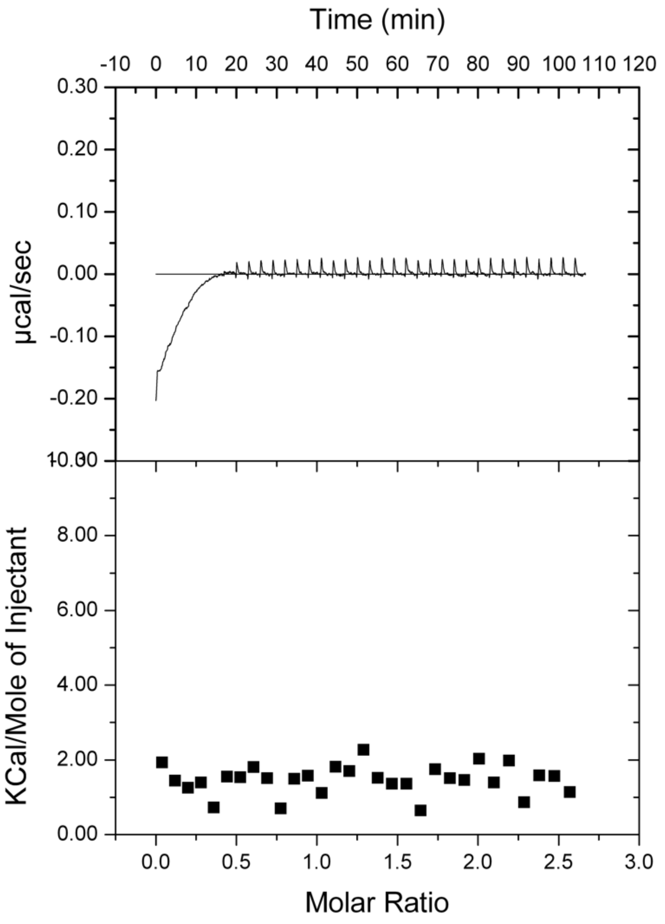

**Author Contributions**

H.L. and L.S.I. conceived the project. G.Z., W.X., Y.-S.T., F.F., D.R.S., H.L. and L.S.I. designed experiments. G.Z. J.I., H.S. and W.X. performed the experiments and analysed the data. G.Z., D.R.S., H.L., P.J.E.R. and L.S.I. wrote the manuscript.
